# Cas4 nucleases can effect specific integration of CRISPR spacers

**DOI:** 10.1101/486738

**Authors:** Zhufeng Zhang, Saifu Pan, Tao Liu, Yingjun Li, Nan Peng

## Abstract

CRISPR-Cas systems incorporate short DNA fragments from invasive genetic elements into host CRISPR arrays in order to generate host immunity.Recently, we demonstrated that the Csa3a regulator protein triggers CCN PAM-dependent CRISPR spacer acquisition in the subtype I-A CRISPR-Cas system of *Sulfolobus islandicus*. However, the mechanisms underlying specific protospacer selection and spacer insertion remained unclear. Here, we demonstrate that two Cas4 family proteins (Cas4 and Csa1) have essential roles (a) in recognizing the 5’ PAM and 3’ nucleotide motif of protospacers and (b) in determining both the spacer length and its orientation. Furthermore, we identify uncovered amino acid residues of the Cas4 proteins that facilitate these functions. Overexpression of the Cas4 and Csa1 proteins, and also of an archaeal virus-encoded Cas4 protein, resulted in strongly reduced adaptation efficiency and the formers yielded a high incidence of PAM-dependent atypical spacer integration, or of PAM-independent spacer integration. We further demonstrated that, in the plasmid challenging experiments, overexpressed Cas4-mediated defective spacer acquisition, in turn, potentially enabled targeted DNA to escape subtype I-A CRISPR-Cas interference. In summary, these results define the specific involvement of diverse Cas4 proteins in *in vivo* CRISPR spacer acquisition. Furthermore, we provide support for an anti-CRISPR role for virus-encoded Cas4 proteins that involves compromising CRISPR-Cas interference activity by hindering spacer acquisition.

**Importance:** The Cas4 family endonuclease is an essential component of the adaptation module in many variants of CRISPR-Cas adaptive immunity systems. The Crenarchaeota *Sulfolobus islandicus* REY15A encodes two *cas4* genes *(cas4* and *csa1)* linked to the CRISPR arrays. Here, we demonstrate that Cas4 and Csa1 are essential to CRISPR spacer acquisition in this organism. Both proteins specify the upstream and downstream conserved nucleotide motifs of the protospacers and define the spacer length and orientation in the acquisition process. Conserved amino acid residues, in addition to the recently reported, were identified to be important for above functions. More importantly, overexpression of the *Sulfolobus* viral Cas4 abolished spacer acquisition, providing support for an anti-CRISPR role for virus-encoded Cas4 proteins that inhibit spacer acquisition.

## INTRODUCTION

The clustered regularly interspaced short palindromic repeats (CRISPRs) and the CRISPR-associated (Cas) proteins generate a diversity of immune systems in most archaea and many bacteria that target invasive viruses and plasmids (1, 2). These diverse systems have been classified into two major classes and at least 6 basic types (Types I to VI) that are further divided into multiple subtypes (3). Spacer acquisition into CRISPR arrays constitutes the first stage of the immune reaction (4); however, the molecular mechanisms involved in this process are still only partially understood. Short DNA fragments of similar length are excised from the invasive genetic element and integrated into the CRISPR loci facilitated by the conserved core proteins Cas1 and Cas2 (1). The first successful demonstration of spacer acquisition under laboratory conditions was obtained for the *Streptococcus thermophilus* subtype II-A system (5) and more recent studies have focused mainly on Type I systems (6) including the subtype I-A systems of *Sulfolobus solfataricus* and *Sulfolobus islandicus* (7–9), the subtype I-B system of *Haloarcula hispanica* (10), the subtype I-E system of *Escherichia coli* (11–13), and the subtype I-F systems of *Pseudomonas aeruginosa* (14) and *Pectobacterium atrosepticum* (15).

Spacer acquisition occurs by two related but differing mechanisms via *de novo* acquisition and primed acquisition pathways. The former requires Cas1 and Cas2 in the subtype I-E system of *E. coli* (12, 16–18), Cas1, Cas2, Csa1, and Cas4 in the subtype I-A system of *S. islandicus* (19), and Cas1, Cas2, Cas9, and Csn2 in *S. thermophilus* subtype II-A systems (20, 21). In contrast, primed acquisition involves activation by a pre-existing spacer in a CRISPR locus that matches the targeted genetic element and this then triggers spacer acquisition for different Type I CRISPR–Cas systems (15–17, 22, 23).

Protospacer-adjacent motifs (PAM) in the DNA of invasive genetic elements determine the acquisition efficiency and integration specificity during the spacer acquisition process. This motif has been shown to direct spacer acquisition for several archaea and bacteria (7, 9, 10, 12, 13). Moreover, the protospacer 3’-terminal motif is important for acquisition efficiency in the subtype I-E system of *E. coli* (13). Different subtypes adapt spacers from invasive genetic elements with a range of lengths. For example, most spacers (95%) in subtype I-E CRISPR arrays of *E. coli* are 32 bp (24) and that is likely to reflect the structure of the substrate-binding Cas1-Cas2 complex (25). But for other systems, including those of subtypes I–A and II–A (26, 27) spacer lengths vary and, moreover, they require proteins additional to Cas1 and Cas2 for specific adaptation (19–21). Most of these additional proteins belong to the Cas4 family and carry nuclease activities. Recently, we showed that two Cas4 family proteins, Cas4 and Csa1, are essential for CRISPR adaptation in *S. islandicus* (19) and more recent studies have shown that the Cas4 nucleases can facilitate PAM recognition and influence spacer lengths and adaptation efficiency (28–30). In particular, the two Cas4 proteins (Cas4-1 and Cas4-2) in *Pyrococcus furiosus* subtype I-A system have been observed distinct activities: Cas4-1 specifies the upstream PAM while Cas4-2 specifies the conserved downstream motif (30).

*S. islandicus* REY15A has proven to be a successful model organism for studying crenarchaeal CRISPR–Cas systems. It carries one subtype I-A and two functionally different subtype III-B interference modules that share an adaptation module encoded adjacent to the subtype I-A interference module (31) and, importantly, a comprehensive array of genetic tools has been developed for this strain (26, 32). Recently, we demonstrated that Csa3a, the activator protein of the subtype I-A CRISPR-Cas immune response in *S. islandicus*, couples transcriptional activation of spacer acquisition and DNA repair genes and, moreover, it enhances CRISPR-Cas interference of invading genetic elements by activating transcription of CRISPR arrays (9, 19). We also demonstrated that the Cas4 family protein genes, *cas4* and *csa1*, were essential for spacer acquisition (19). However, the mechanisms involved require further study including determining the essential amino acid residues of the different Cas4 proteins. Moreover, the functions of virus-encoded Cas4 family proteins remain to be determined.

In the present study, we focus on the functions of the Cas4 family proteins, Cas4 and Csa1 in CRISPR spacer acquisition in *S. islandicus*. We have found new *in vivo* characteristics of *Sulfolobus* Cas4 compared with previous *in vitro* works on *Sulfolobus* Cas4 and with Cas4 proteins of other organisms. Moreover, we provide evidence that *Sulfolobus* virus-encoded Cas4 proteins may play an important role in reducing the efficiency of host CRISPR-Cas interference.

## RESULTS

### Cas4 family proteins are divided into different subtypes

Many CRISPR-Cas adaptation systems require proteins additional to Cas1 and Cas2 and, in particular, Cas4 family proteins that are encoded widely in archaeal and bacterial genomes and in their viruses. A phylogenetic tree is presented (Fig. 1) derived from alignments of 28 Cas4 family proteins that include 21 genome-encoded Cas4 proteins from six archaeal genomes and seven archaeal viral Cas4-like proteins. Here, Cas4_Sso1392 does not carry the four conserved cysteine residues (4-C motif) which forms a Fe-S cluster. The sequences group into 4 subtypes, CRISPR-related Cas4, CRISPR-related Csa1, solo Cas4 and viral Cas4-like proteins, where the genes of solo Cas4 and viral Cas4 proteins are not linked to CRISPR loci (Fig. 1).

**Figure 1.**
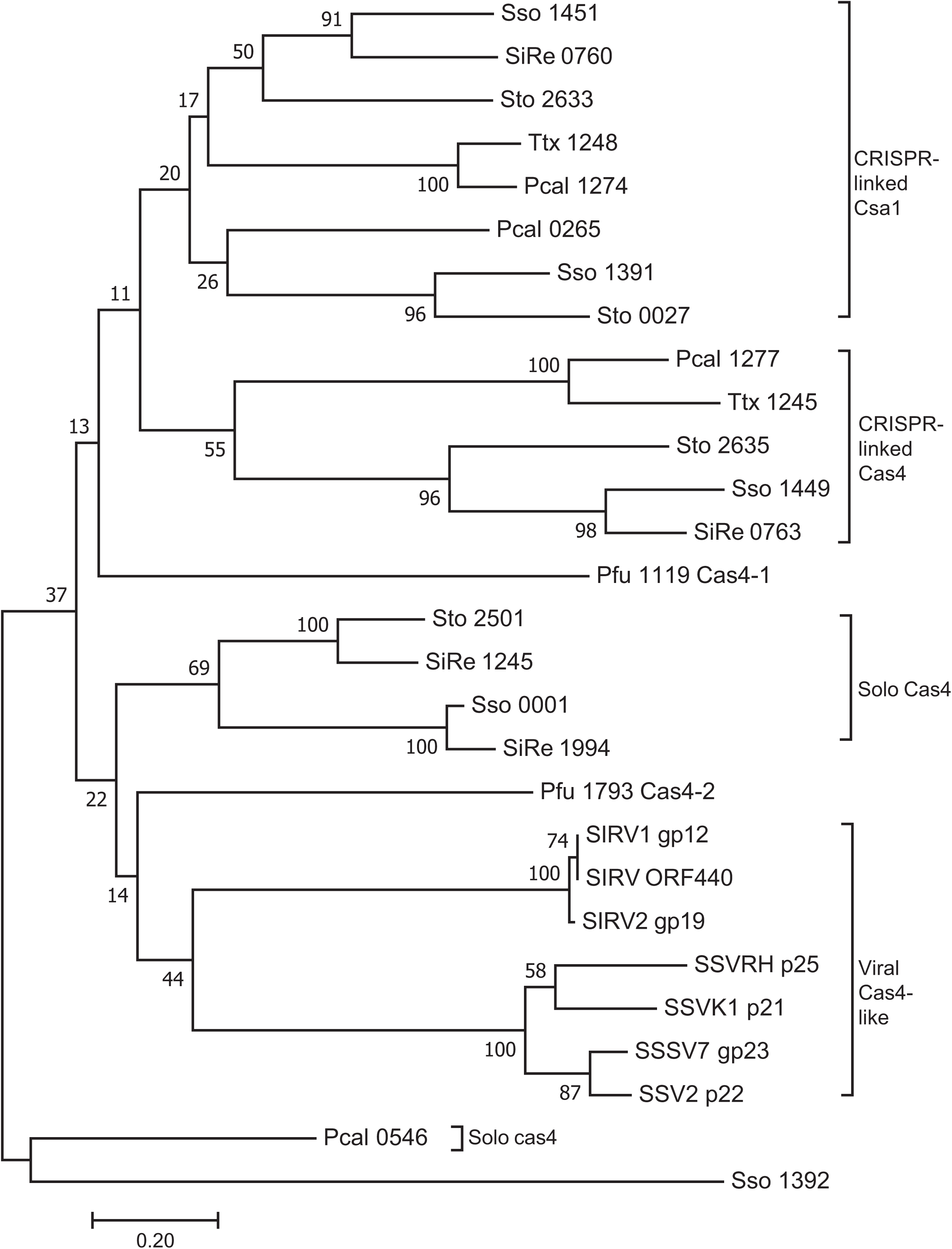
Unrooted bootstrapped phylogenetic tree showing a subset of archaea and their viruses-encoded Cas4 family proteins. Each protein is represented by a species code followed by the gene number (Sso, *S. solfataricus*; SiRe, *S. islandicus* REY15A; Pfu, *P. furiosus*; Sto, *S. tokodaii*; Ttx, *T. tenax*; Pcal, *Pyrobaculum calidifontis*; SIRV, *Sulfolobus islandicus* rod-shaped virus; SSV, *Sulfolobus* Spindle-shaped Viruses; SSSV7, *Sulfolobus* Spindle-Shaped Virus 7;). This neighbour-joining tree was generated from a T-coffee alignment of the proteins using Mega7, with pairwise distances between sequences uncorrected. The bootstrap values shown at each node represent the percentage of all trees (10000 total) agreeing with this topology. The subgroup of CRISPR-associated Csa1, CRISPR-associated Cas4, solo Cas4, viral Cas4 like proteins are indicated.

*Sulfolobus* species generally encode multiple Cas4 proteins (Fig. 1). For example, in the crenarchaeon *S. islandicus* REY15A, *cas4* and *csa1* genes neighbour CRISPR arrays while two other *cas4* genes are not linked to CRISPR arrays (Fig. 1). The euryarchaeon *Pyrococcus furiosus* JFW02 carries one CRISPR-linked *cas4-1* gene and a second solo *cas4-2* gene (Fig. 1). However, although the Cas4-1 (Pfu 1119)-encoding gene adjoins a CRISPR array, it diverges strongly from the *Sulfolobus* CRISPR linked Csa1 or Cas4 (Fig. 1). Similarly, *P. furiosus* Cas4-2 (Pfu 1994) falls between solo Cas4 and viral Cas4-like proteins (Fig. 1). Several *Sulfolobus* rudiviruses and fuselloviruses encode Cas4 proteins and their genes cluster in the phylogenetic tree (Fig. 1). This raises two questions: (1) what are the functions of CRISPR-linked Cas4 and Csa1 and the solo Cas4 proteins in the *Sulfolobus* genome and (2) what effects, if any, do the viral Cas4 proteins have on the CRISPR-Cas reponse of the infected host.

### Cas4 proteins define spacer origin and recognise the PAM and the 3’ conserved DNA motifs

*S. islandicus* REY15A encodes one spacer acquisition module, constituting Cas1, Cas2, Cas4, and Csa1 (Fig. 2A). The downstream gene, *csa3a*, encodes a transcriptional regulator that activates expression of the adaptation module (9). This, in turn, triggers CCN PAM-dependent *de novo* spacer acquisition (9, 19). Earlier, we demonstrated that deletion of the *cas4* and *csa1* genes eliminated the capacity for CRISPR spacer acquisition as judged by the inability of the deletion strain to produce larger PCR products in the CRISPR leader-proximal region (19).

**Figure 2.**
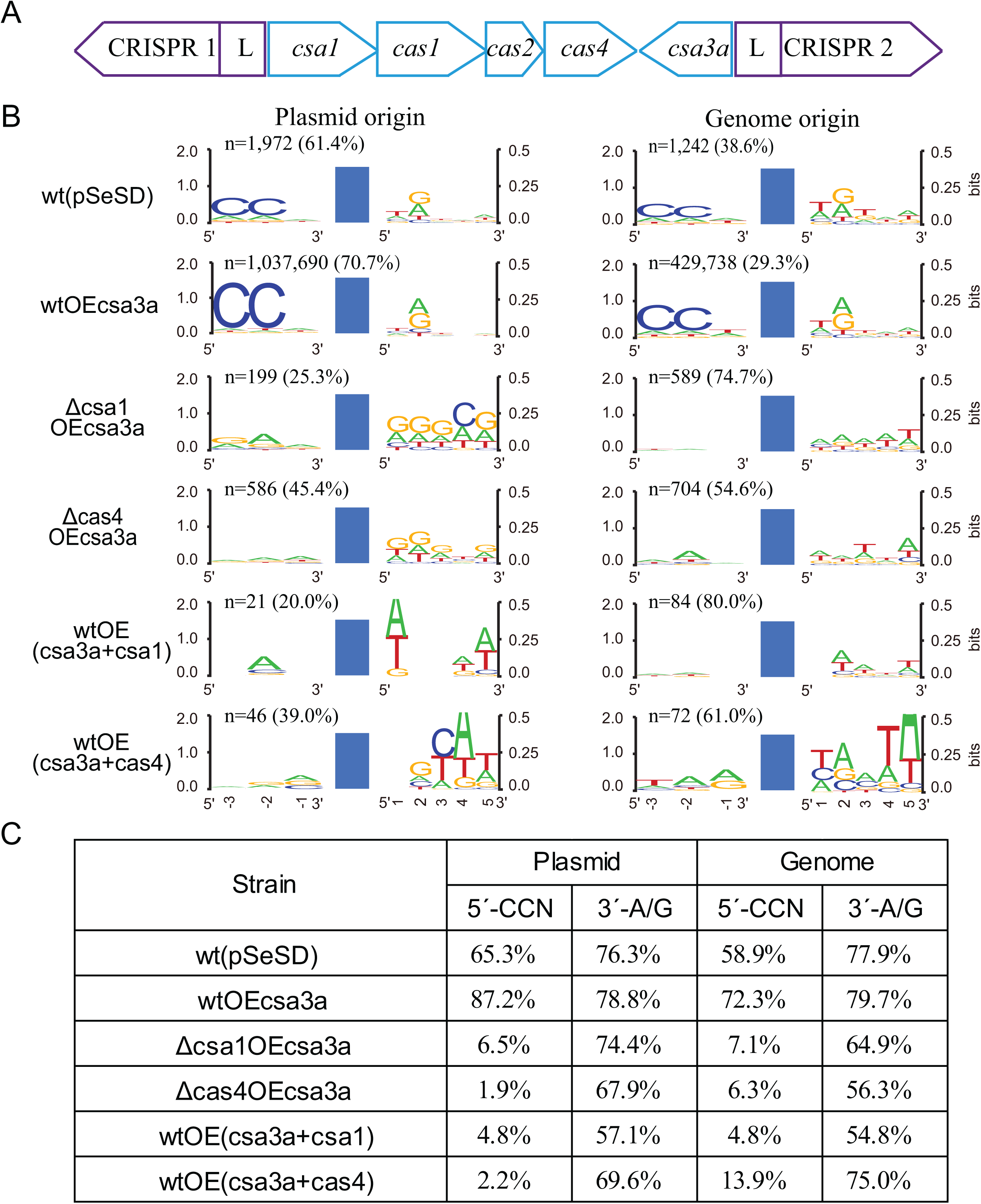
Effects of Cas4 proteins on spacer origin, PAM and 3’ conserved motif. **A**, Organization of the CRISPR-Cas adaptation module of *S. islandicus* REY15A with two CRISPR arrays. **B**, Weblogo analysis of the conserved motifs upstream and downstream of the protospacers detected in different *Sulfolobus* transformants by high-throughput sequencing data. Reads of the new spacers (n) and the percentage of plasmid- or genome-derived spacers are indicated. wt(pSeSD): wild-type cells carrying empty vector; wtOEcsa3a: wt cells overexpressing *csa3a* activator gene; Δcsa1OEcsa3a: *csa1* deletion strain overexpressing *csa3a;* Δ*cas4*OEcsa3a: *cas4* deletion strain overexpressing *csa3a;* wtOE(csa3a+csa1): wt cells overexpressing both *csa3a* and *csa1*; wtOE(csa3a+cas4): wt cells overexpressing both *csa3a* and *cas4*. Blue columns: protospacers. **C**, percentage of the new spacers of plasmid or genome origin carrying CCN or A/G motifs from the different genetically modified strains.

In this study, we examine the effects of deletion and overexpression of Cas4 and Csa1 on spacer acquisition by high-throughput sequencing, and analysis, of newly acquired spacers. The high throughput data were summarized in supplementary Table 1. Consistent with the earlier work (9,19), the high-throughput sequencing data revealed that the control strain, carrying only the empty vector pSeSD, yielded relatively few new spacers while many were observed in wt cells with *csa3a* overexpression (Fig. 2B). Moreover, in both strains, most new spacers derived from the expression vector and few were from host genome DNA (Fig. 2B). Next, we examined the effects of Cas4 and Csa1 on spacer acquisition by testing a deletion strain of *cas4*, and of *csa1* (Fig. 2B). Very few new spacers were detected for either deletion strain which indicated that Cas4 and Csa1 were essential for specific spacer acquisition. In addition, we demonstrated that enhancing expression of Cas4, and Csa1 also, strongly reduced spacer acquisition (Fig. 2B), probably due to excess Cas4 or Csa1 proteins forms large complex such as a decameric toroid as reported previously (33) to disorder the adaptation complex. Furthermore, when we examined the origins of the spacers we found that the relatively few spacers that were acquired in the *cas4* and *csa1* deletion or overexpression mutants did show a stronger bias to genomic DNA than to the plasmid (Fig. 2B). This observation also reinforced that the Cas4 proteins were crucial for specific selection of spacers.

Since CRISPR-Cas interference requires PAM recognition, we analyzed the PAM and 3’-nucleotide motif of the cognate protospacers. New spacers in the wt strain carrying the empty vector derived predominantly from protospacers carrying conserved CCN PAM and 3’-A/G motifs (Fig. 2B and C). Moreover, protospacers in the *csa3a* overexpression strain also predominantly carried CCN PAM and the 3’-A/G motifs (Fig. 2B and C), consistent with our earlier results (9). In contrast, the CCN PAM was absent from protospacers of both the deletion, and overexpression, strains of *cas4* and *csa1* (Fig. 2B and C); the occurrence of the 3’-A/G motif at the +2 site was also reduced in these strains (Fig. 2B and C). These results indicated that proteins Cas4 and Csa1 play a role in locating the PAM and 3’-motif during spacer selection. Thus, the absence of the PAM in these strains will prevent CRISPR-Cas interference from the defective spacers. Potentially, new spacers can accumulate from any DNA site in the four strains and this will produce a strong bias to host genomic DNA because of the much higher DNA yields relative to those of the low copy number *csa3a* overexpression plasmid (Fig. 2B). It should be noted that few new spacer reads (~ 100 reads) were identified in the *csa1* or *cas4* overexpression strains (Fig. 2B). PAM sequence from these few reads could showed less conservation. However, new spacers from *csa1* or *cas4* overepxression strains were matched to different locations on the genomic and plasmid DNA, and the ratio between unique new spacer reads and is total new spacer reads 0.60 and 0.23, suggesting identification of non-PAM in this these strains may reflect the truth.

### Novel amino acid residues essential for Cas4 protein functions

Cas4 family proteins carry a few highly conserved amino acid residues (34) (Fig. 3A) and we examined whether they were involved in specific protospacer selection. Firstly, we analysed the amino acid residues in the RecB motif of the 28 archaeal genome-encoded or archaeal virus-encoded Cas4 proteins (Fig. 3B). We identified a conserved Asp residue in *S. islandicus* Cas4 (D60 in SiRe_0763) and the CRISPR-linked Csa1 proteins, and two conserved Trp/Leu residues in CRISPR-linked Cas4 proteins (YL100/101 in SiRe_0763) (Fig. 3B). Therefore, we mutated these amino acids separately with a view to establish whether they were essential for specific spacer acquisition.

**Figure 3.**
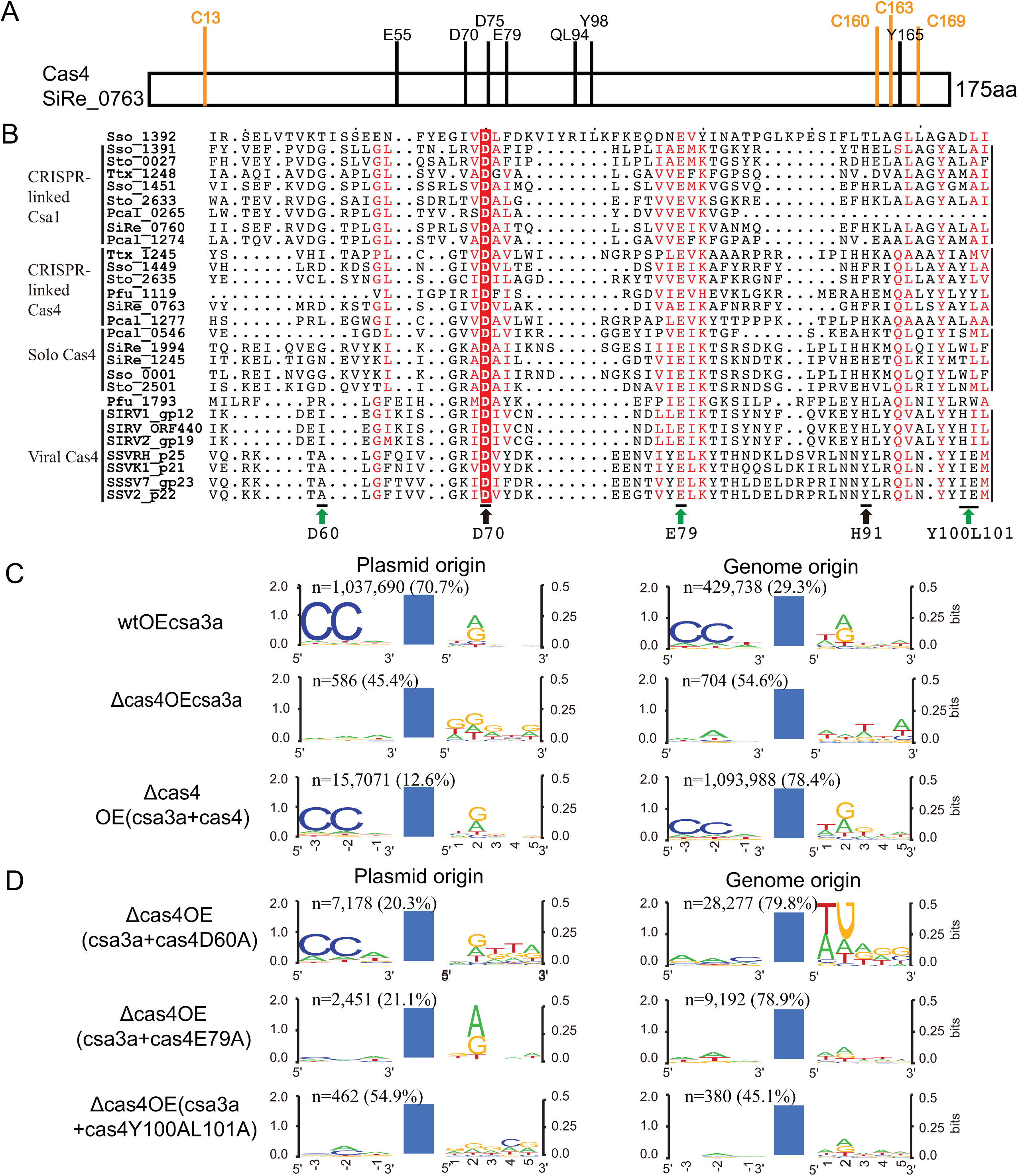
Identification of amino acid residues of Cas4 nucleases that influence spacer origin, PAM and 3’ conserved motif recognition. **A**, diagram showing conserved amino acid residues in *S. islandicus* Cas4 protein (SiRe_0763). **B**, Aligment of the RecB domain of Cas4 family proteins. Green arrows indicate the mutations in this study; black arrows indicate mutations analysed earlier (30). Subgroups of the Cas4 family proteins are indicated. **C** and **D**, Weblogo analysis of the conserved motifs upstream or downstream of protospacers detected in different *Sulfolobus* transformants from high-throughput sequencing data. Reads of the new spacers and the percentage of plasmid- or genome-derived spacers are indicated. Blue columns: protospacers. wtOEcsa3a: wt cells overexpressing the *csa3a* activator gene; Δ*cas4*OE3a: *cas4* deletion strain overexpressing *csa3a;* Δ*cas4*OE(csa3a+cas4): *cas4* deletion strain overexpressing *csa3a+cas4;* Δ*cas4*OE(csa3a+cas4D60A): *cas4* deletion strain overexpressing *csa3a*+*cas4*D60A; Δ*cas4*OE(csa3a+cas4E79A): *cas4* deletion strain overexpressing *csa3a+cas4*E79A; Δ*cas4*OE(csa3a+cas4Y100AL101A): *cas4* deletion strain overexpressing csa3a+cas4Y100AL101A.

The *cas4* and mutant genes cas4D60A, cas4E79A and cas4Y100A/L101A were cloned downstream of the *csa3a* gene on plasmid pSeSD, and expression of *csa3a, cas4* and the *cas4* mutants were under control of the *araS* promoter (35). These plasmids were then transformed into the *cas4* deletion cells to yield complementary strains. Whereas transformation of p(csa3a+cas4) reactivated highly efficient spacer acquisition, albeit with a strong bias to genomic DNA (Fig. 3C), transformation with p(csa3a+cas4D60A) generated a low level of specific spacer acquisition, again with a bias to genomic DNA, while transformation with the other two p(csa3a+cas4 mutant) plasmids failed to restore specific spacer acquisition (Fig. 3D). We infer from the latter result that Cas4 positions E79 and Y100/L101 are critical for selection of the CCN PAM in protospacers.

### Cas4 facilitates specific spacer integration

As demonstrated the CCN PAM was not detected in most protospacers from the *cas4* and *csa1* deletion or overexpression strains or in those deriving from genomic DNA, as indicated in Fig. 2B, Fig. 3C and D). These results demonstrate that at suboptimal-levels of Cas4 unspecific spacer integration occurred. This result provided an explanation for the majority of spacers deriving from the much more prevalent host genomic DNA. Complementing *cas4* in the *cas4* deletion mutant strain that overexpressed *csa3a* restored the selection of protospacers with conserved 5’-CCN PAMs (Fig. 4). Complementing the *cas4* deletion mutant with cas4D60A, cas4E79A and cas4Y100AL101A completely or partially restored selection of plasmid DNA protospacers carrying CCN PAMs (Fig. 4A) but not for those deriving from genomic DNA (Fig. 4 B). However, CCN PAM-dependent inverted integration and slipped integration (atypical CCN) accounted for a large proportion in cas4D60A complementing strain (Fig. 4B), suggesting cells with these genomic protospacers survived from CRISPR interference guided by the flipped and slipped integrated spacers. These results indicate that amino acid residues D60A, E79 and YL100/101 are important for selection of specific protospacers in *Sulfolobus*. Particularly, complementing *cas4E79A* caused half of the genomic protospacers with no CCN PAM at either end (Fig. 4B), suggesting it was extremely important for CCN PAM selection.

**Figure 4.**
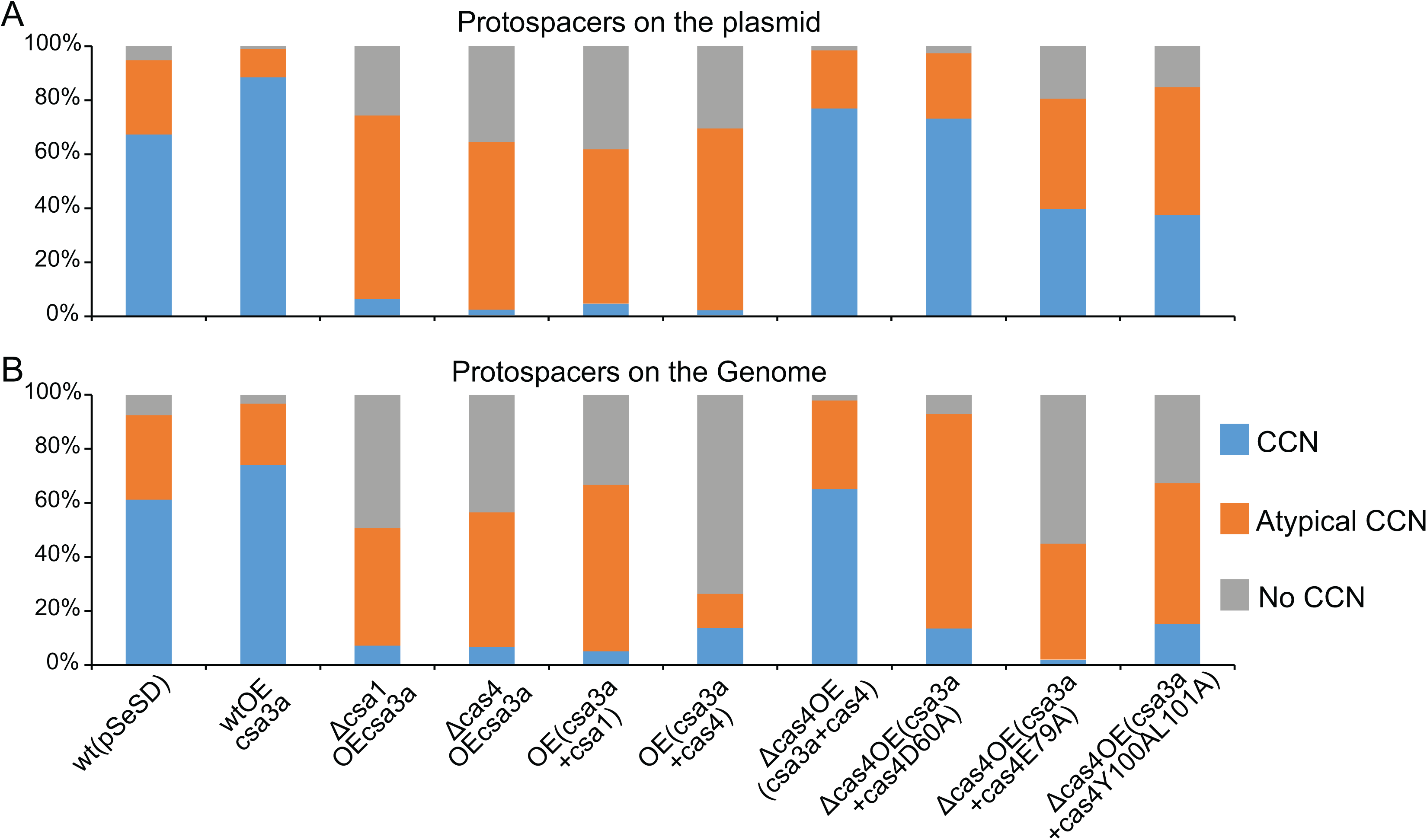
Percentage of protospacers on the plasmid (A) and on genomic DNA (B) that carried conserved CCN, atypical CCN and no CCN motifs. Blue (CCN): CCN motif at the 5’ adjacent of protospacers; Orange (Atypical CCN): the spacers with flipped, slipped, flipped-slipped integration or with CDN or DCD motif (a single conserved C at the −2 or −3 site of the 5’ adjacent region); Grey (No CCN): the cognate protospacers with no CCN motif at either end. All cognate protospacers were detected in high-throughput sequencing data.

### Cas4 nuclease defines spacer length

Spacer lengths vary in *Sulfolobus* species although they fall mainly in the range 39–41 bp (26). Most spacers acquired in wild-type *S. islandicus* strains, carrying pSeSD or *csa3a* overexpression vectors, were 40–41 bp (Fig. 5) similar to the lengths of spacers present in the host CRISPR arrays. However, deletion of *cas4* or *csa1* not only strongly reduced spacer acquisition efficiency, as measured by the n value (Fig. 2B and 3C), but it also yielded, on average, slightly shorter spacer lengths (Fig. 5). Overexpression of *csa1* and *cas4* also produced altered length distributions (Fig. 5). Complementing *cas4* in the *cas4* deletion strain did restore spacer lengths to the normal host size distribution but whereas complementing cas4Y100AL101A did not alter average spacer lengths, while complementing cas4D60A and E79A produced increased length distributions (Fig. 5). These results reinforce further that Cas4 is important for producing specific spacer acquisition.

**Figure 5.**
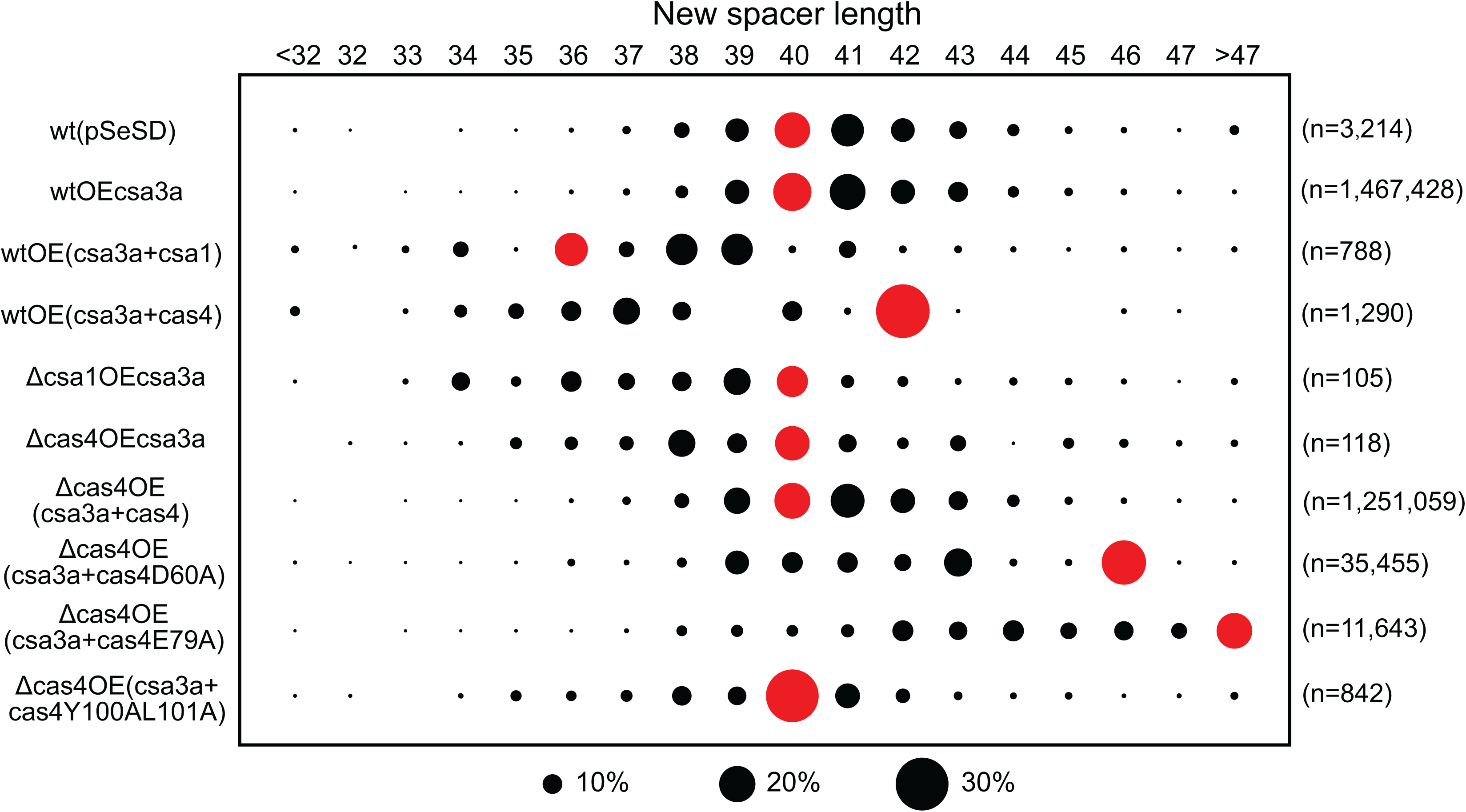
Cas4 nucleases define spacer length. Spacer lengths are shown in the upper row. Strains used were shown in left panel. Reads of the new spacers were indicated at the right panel. Filled bubbles showed the percentage of protospacers, and the red bubbles indicated the spacer length of majority.

### Flipped and slipped spacers inactivate DNA interference of the subtype I-A system

In order to demonstrate that flipped spacers abolish subtype I-A CRISPR-Cas interference activity, we designed a plasmid challenge experiment that is illustrated in Fig. 6A. The DNA sequence of spacer 10 from CRISPR locus 2 of *S. islandicus* REY15A was cloned into the *Sulfolobus-E. coli* shuttle vector pSeSD under control of the *araS* promoter (35) with different 5’- or 3’-motifs to either activate, or escape from, subtype I-A and III-B DNA interference activities (Fig. 6A). These challenging plasmids, and the control expression vector pSeSD, were transformed into *S. islandicus* E233S and transformation efficiencies were estimated. If the plasmids were targeted by the subtype I-A and III-B interference complexes, guided by crRNA from spacer 10, the transformation efficiency would be strongly reduced.

**Figure 6.**
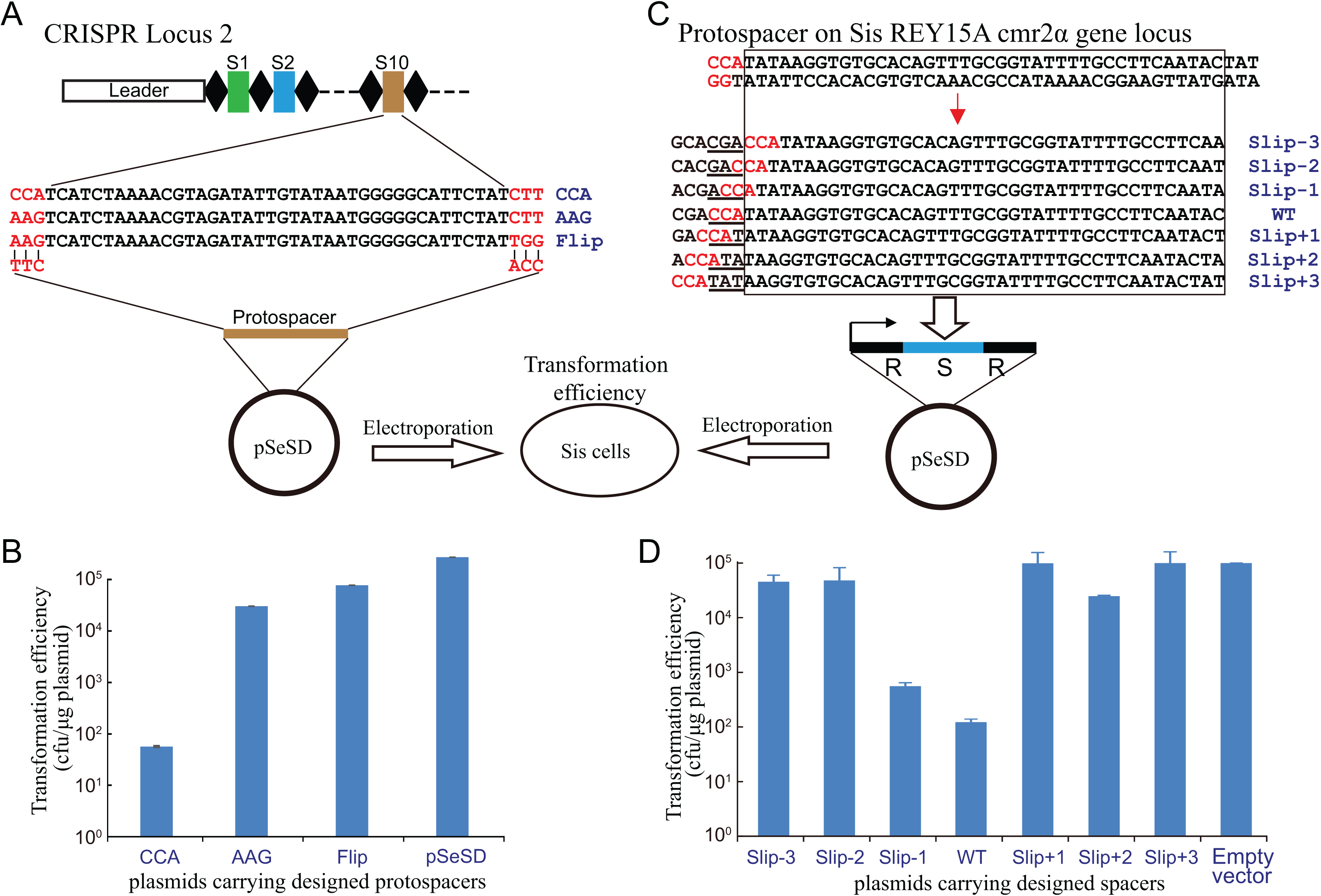
Influence of flipped and slipped spacers on DNA interference activity of the subtype I-A system. A, The DNA sequence of spacer 10 in CRISPR locus 2 was selected and inserted into the *Sulfolobus-E. coli* shuttle vector pSeSD along with the indicated 5’-end and 3’-end adjacent sequences. B, Transformation efficiencies of plasmids carrying flipped and control spacers electroporated into *S. islandicus* E233S. C, A protospacer with the CCA PAM sequence matching the template strand of the *cmr2a* gene was selected, and the cognate spacer and its slipped derivatives were cloned into the site between two repeats in the pSeSD plasmid to generate the mini-CRISPR expression vectors. D, Transformation efficiencies of plasmids carrying the mini-CRISPR cassettes electroporated into *S. islandicus* E233S.

The plasmid carrying the protospacer matching spacer 10 with a 5’-CCA PAM showed a very low transformation efficiency compared with that of the control plasmid, consistent with strong subtype I-A DNA interference having occurred (Fig. 6B). Another control plasmid carrying the same protospacer with a 5’-AAG motif showed a relatively low transformation efficiency compared with pSeSD plasmid but a much higher transformation efficiency compared with the plasmid carrying the CCA motif (Fig. 6B). This indicated that the 5’-AAG motif inhibited subtype I-A interference. The third plasmid carrying the spacer with protospacer 5’-AGG and 3’-TGG sequences mimics the protospacer of the flipped spacers inserted into the CRISPR array (Fig. 6A). High ‘flip’ plasmid transformation efficiency, similar to that of the AAG plasmid (Fig. 6B) would indicate that flipped spacers had integrated into CRISPR arrays and were inactive in subtype I-A.

Slipped spacer integration can also impede subtype I-A DNA interference because the crRNAs will fail to recognise the CCN PAM sequence. To test this experimentally, we selected a protospacer from the template strand of the *S. islandicus* REY15A *cmr2α* gene and generated different mutants carrying single site mutations (Fig. 6C). The altered spacers were cloned into the *Sulfolobus* expression vector pSeSD (35) to generate the plasmid-borne mini-CRISPRs under the control of the arabinose-inducible promoter (Fig. 6C) (32, 36). These challenging plasmids were then transformed into *S. islandicus* E233S cells and transformation efficiencies were estimated. If the plasmid-borne spacer crRNA causes interference at *cmr2α*, then low transformation efficiency will be observed, and *vice versa*. Since the designed spacer is from the *cmr2α* template strand, the crRNA will not base-pair with the mRNA and, therefore, will not trigger the subtype III-B DNA interference activity. The transformation efficiency of pSeSD carrying the wild-type protospacer was very low compared with that of the empty vector, consistent with strong subtype I-A interference of *cmr2α* (Fig. 6D). The plasmid carrying the Slip-1 spacer (corresponding to the protospacer with the NCC motif) showed a transformation efficiency that was lower than the empty vector control but higher than the wild-type spacer plasmid, indicating that moderate plasmid interference had occurred (Fig. 6D). This reflects that a 5-ACC motif of the protospacer can mediate subtype I-A interference of the target DNA with reduced efficiency (Fig. 6D). No interference was observed at *cmr2α* by the other Slip spacers, except for very weak interference by the Slip+2 spacer (Fig. 6D). We conclude that the slipped spacers strongly reduced or eliminated subtype I-A CRISPR-Cas interference.

### A virus-encoded Cas4 protein hinders CRISPR acquisition

The *cas4* genes are present in most known viruses of the Sulfolobales (37). *Sulfolobus* spindle-shaped viruses (SSV) encode Cas4 that cluster with a group of Cas4 proteins associated with type I CRISPR-Cas systems of the Thermococcales, and a few Cas4 proteins encoded by Icelandic rudiviruses (eg. SIRV in Fig. 1) also form a potential branch in this family (37). However, even the widespread of *cas4* genes in viral genomes, the functions of viral Cas4 on CRISPR spacer acquisition or on virus-hots interaction are still unknown. In this study, the *cas4* gene of SSV2 was cloned into the *csa3a* overexpression plasmid and transformed into *S. islandicus* in order to study its effect on CRISPR spacer acquisition. The positive control strain overexpressing *csa3a* (wtOE3a) acquired new spacers, as evidenced by the expanded PCR bands deriving from the leader-proximal CRISPR region (Fig. 7A). In contrast, two strains from single colonies overexpressing both *csa3a* and SSV2 *cas4* failed to yield detectable new spacers (Fig. 7A). PCR products from the leader-proximal CRISPR regions of each of the above strains were subsequently sequenced and and the results demonstrated that expression of SSV2 Cas4 produced strongly reduced CRISPR spacer acquisition efficiency (<1%) (Fig. 7B). We analysed the 5’- and 3’-ends of the corresponding protospacers from the viral *cas4* expression strains, and found that whereas most protospacers of plasmid origin carried a conserved CCN PAMs (70.7%), those of genome origin exhibited a lower level of 5’-CCN PAM (29.3%) (Fig. 7B). This result suggested the new spacers were active activating CRISPR subtype I-A interference and that surviving cells carried protospacers lacking PAM motifs. In addition, overexpression of SSV2 *cas4* led to reduced conservation of the protospacer 3’-A/G motif (Fig. 7B). The length distribution of the adapted spacers from both viral *cas4* expression strains was similar to the wt cells (Fig. 7C), indicating viral Cas4 had no effect on spacer length. In summary, expression of SSV2 Cas4 protein strongly reduced spacer acquisition efficiency and reduced the conservation of 3’- A/G motif (Fig. 7A and B) but did not lead to changes in the 5’-CCN PAM or the spacer length (Fig. 7B and C).

**Figure 7.**
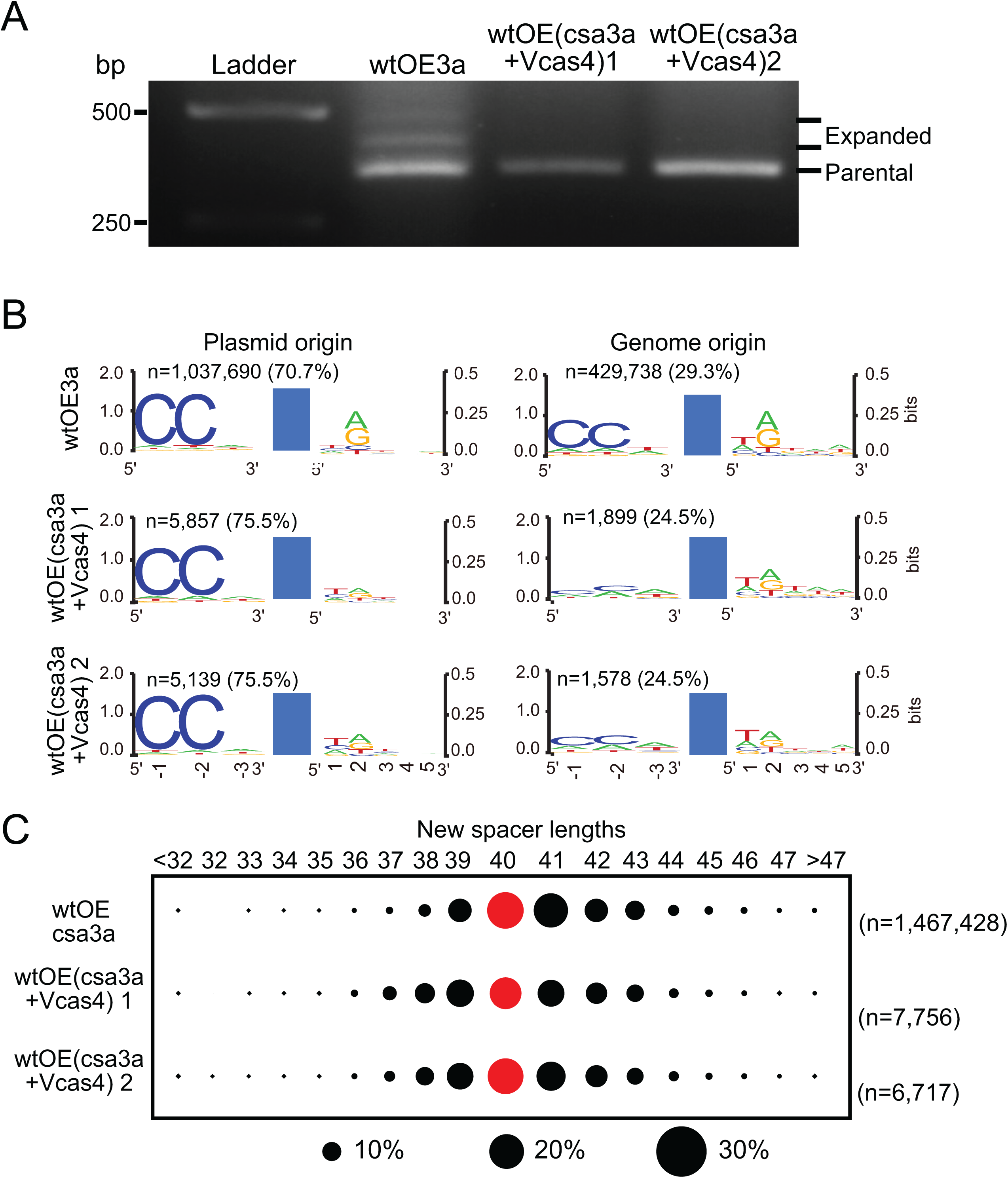
The effect of a viral Cas4 protein on CRISPR spacer acquisition in *Sulfolobus*. **A**, PCR analysis of the leader proximal region of CRISPR locus 1 in *S. islandicus* transformants. OEcsa3a: wt cells overexpressing *csa3a* activator gene; OE(csa3a+Vcas4) 1 and OE(csa3a+Vcas4) 2: cells of single colonies 1 and 2 of wt cells overexpressing both *csa3a* and SSV2 *cas4*. **B**, Analysis of the conserved motifs upstream and downstream of the protospacers detected in different *Sulfolobus* transformants by high-throughput sequencing data analyses using Weblogo. **C**, New spacer lengths are shown in the upper row. Strains used are indicated in the left panel. Filled bubbles showed the percentage of protospacers, and the red bubbles indicated the spacer length of majority. Reads of the new spacers (n) and the percentage of plasmid- or genome-derived spacers are indicated.

## DISCUSSION

### Conserved amino acid residues define Cas4 functions

Cas4 family proteins occur widely within CRISPR-Cas adaptation modules of subtype I-A, I-B, I-C, I-D, I-U, II-B, and type V systems (2) and they include Csa1 of *Sulfolobus* subtype I-A systems and Csn2 of the subtype II-A systems of *S. thermophiles* (20) and *S. pyogenes* (21). High resolution crystal structures of Cas4 proteins from *S. solfataricus* (33) and *P. calidifontis* (38) indicate that they carry two domains, an N-terminal RecB-like nuclease domain and a C-terminal domain containing an Fe-S cluster coordinated by four conserved cysteine residues. Moreover, biochemical studies have shown that Cas4 can exhibit 5’ to 3’ and 3’ to 5’ DNA exonuclease activity, as well as ATP independent DNA unwinding activity (33, 34, 38) which suggests that they produce single-strand DNA overhangs that are potential intermediates for insertion of new CRISPR spacers. Previously, we have demonstrated that both Cas4 and Csa1 proteins are essential for *de novo* spacer acquisition (19). Recently, an *in vitro* study has demonstrated that Cas4 of *Bacillus halodurans* subtype I-C system interacts tightly with Cas1 integrase and processes double-stranded substrates with long 3’ overhangs through site-specific cleavage (29). Cas4 recognizes PAM sequences within the prespacers and prevents integration of unprocessed prespacers, ensuring correct integration (29). An *in vivo* study reveals that Cas4 of subtype I-D system from *Synechocystis* 6803 pSYSA megaplasmid, in addition to Cas1 and Cas2, facilitates recognition of PAM and defines the spacer length in a heterogenous host, *E. coli* (28). Although one conserved amino acid residue of Cas4 protein from *Synechocystis* subtype I-D system has been identified to define PAM and spacer length (28), more works are required to study the functions of Cas4 proteins in their native hosts.

Most recently, the Cas4 functions of *P. furiosus* subtype I-A system have been studied *in vivo* (30). *P. furiosus* JFW02 genome carries two Cas4 genes: *cas4-1* is linked with the CRISPR array, and *cas4-2* is solo. Shiimori et al. observed distinct activities between the two Cas4 proteins, especially in recognition of upstream PAM and downstream conserved DNA motif, respectively, in *P. furiosus* (30). While in our study, we found both Csa1 and Cas4 were crucial for specifying the upstream PAM sequence in *Sulfolobus* (Fig. 2B). Shiimori et al. have also identified one conserved amino acid residues of RecB motif involved in nuclease activity in both Cas4 proteins for defining the spacer length and orientation in *P. furiosus* (30). We experimentally identified three conserved amino acid residue sites of the RecB motif were important for Cas4 function, in addition to the Shiimori and co-workers’ results: the residue D60 conserved in CRISPR-linked Csa1 proteins, Y100L101 conserved in CRISPR-linked Cas4 proteins, and E79 conserved in all Cas4 family proteins (Fig. 3B). We summarize the experimentally studied conserved amino acid residues in RecB motif of Cas4 family proteins involved in different functions in CRISPR spacer acquisition (Fig. 8). D60, identified in *S. islandicus* Cas4, is essential for spacer acquisition and defines spacer length, but it is not responsible for defining 5’-CCN PAM and 3’ nucleotide motif (Fig. 8B). D70, corresponded to *S. solfataricus* Sso0001 is essential to its *in vitro* nuclease activity (34), and D70 corresponds to the residue in Cas4-1 or Cas4-2 of *P. furiosus* determines acquisition efficiency, spacer length and integration orientation, and recognises the 5’- or 3’-protospacer motifs *in vivo* (30)(Fig. 8B). E79, identified in *Sulfolobus* Cas4, influences acquisition efficiency and is important for determining spacer length, integration orientation and the 5’ and 3’ motifs of protospacers (Fig. 8B). However, H91 corresponding to a residue in Cas4-1 or Cas4-2 of *P. furiosus* is unimportant for Cas4 functions (30)(Fig. 8B). YL100/101 identified in this study strongly effects Cas4 functions (Fig. 3D), probably due to that these amino acid residues affect the nuclease activity or the conserved hydrophobic residue L101 is important for Cas4 structure integrity. Similarly, the 4-C cluster of both Cas4-1 and Cas4-2 of *P. furiosus* is essential for Cas4 functions (30), consistent with published findings that the Fe-S cluster is required for the structural integrity of Cas4 proteins (33).

**Figure 8.**
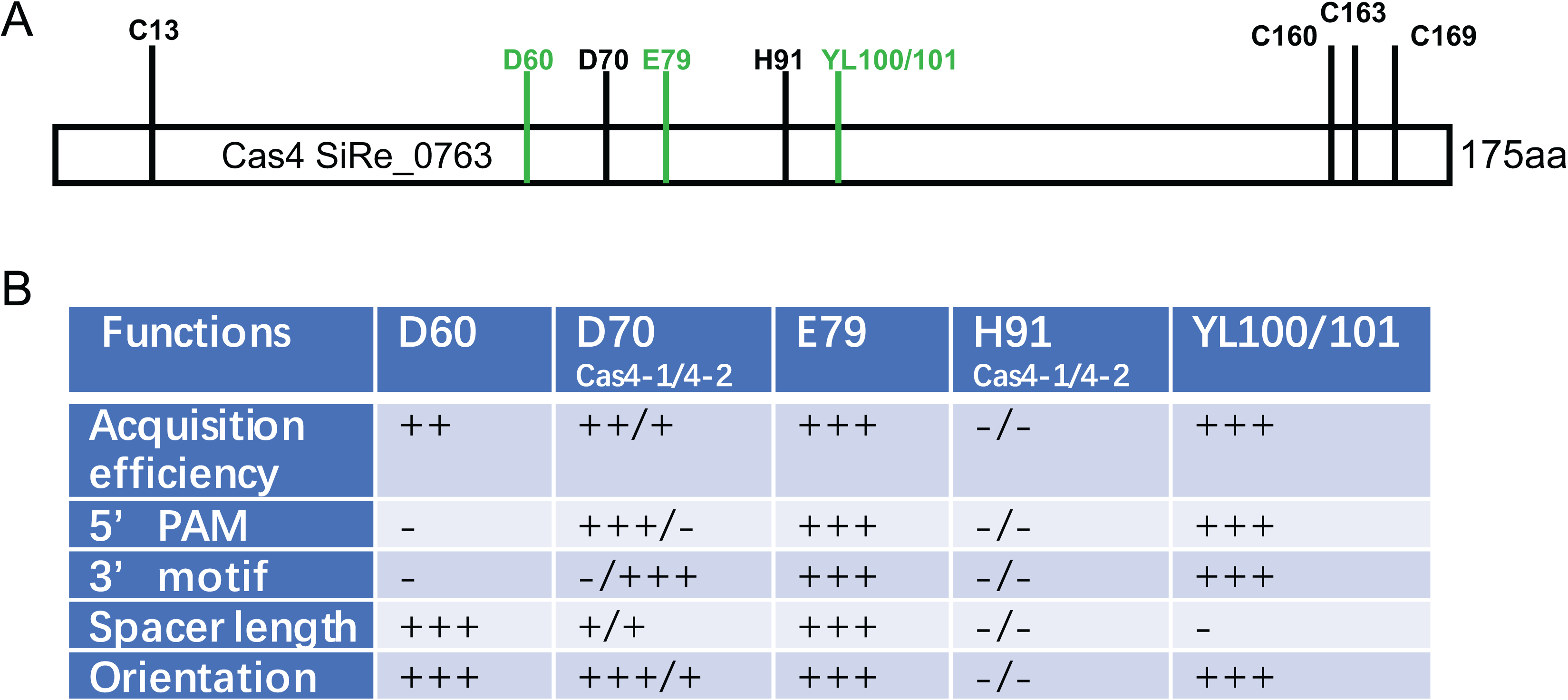
Summary of experimentally studied conserved amino acid residues of Cas4 family proteins involved in the functions of CRISPR spacer acquisition. **A**, Experimentally characterized amino acid residues in Cas4 family proteins, using the Cas4 (SiRe_0763) protein from *S. islandicus* REY15A. Conserved residues represented by black lines and letters were identified by Shiimori et al. (30) and conserved residues represented by green lines and letters were examined in this study. **B** Functions of the experimentally characterized amino acid residues in RecB nuclease motif of Cas4 proteins. Weak to strong influences were indicated by (+) to (+++), and no influence was indicated by (-). D70 and H91 showed their functions in Cas4-1 and Cas4-2 from *P. furiosus* (30).

Csa1 belongs Cas4 family proteins which is specific for subtype I-A CRISPR-Cas systems (2). In this study, we firstly report its functions, similarly to Cas4, in defining PAM, orientation and spacer length (Fig. 2, 4 and 5). Even though Csa1 proteins carry conserved amino acid residues as Cas4 protein do, a long insert is found in Csa1 amino acid sequence, suggesting some different roles may be played by Csa1 proteins.

### A putative anti-CRISPR-Cas mechanism evolved by viruses

In this study, we show that overexpression of Cas4 or Csa1 can trigger a strong reduction in CRISPR-Cas spacer acquisition efficiency (Fig. 2B) and induce a larger fraction of new spacers to be integrated in a flipped or slipped manner, or to be derived from protospacers with no conserved 5’-CNN PAM motif (Fig. 4). Furthermore, *cas4* overexpression yields a much larger ratio of spacers from host genomic DNA relative to plasmid DNA than in the wild-type (Fig. 2B). Therefore, we conclude that even though the host CRISPR loci acquired few new spacers in the *cas4* overexpression strain, the host and plasmid targets evade CRISPR-Cas interference because of defective spacer-protospacer recognition by subtype I-A system. To test this hypothesis, we calculate the transformation efficiencies of plasmids carrying either flipped spacers or mini-CRISPR loci with slipped spacers. Then we demonstrate experimentally that slipped and spacers hindered subtype I-A DNA interference (Fig. 6).

Sub-optimal level of the host encoded Cas4 reduces spacer acquisition efficiency and caused defective acquisition. However, Cas4-like nucleases are frequently encoded by archaeal viruses (39, 40) and by phages (41) and occasionally by transposable elements (42). Moreover, a Cas4-like protein from the rudivirus SIRV2 is shown to possess both 5’- to 3’-exonuclease and endonuclease activities *in vitro* (39, 43), suggesting that the viruses encode functional Cas4 nucleases that could participate in CRISPR spacer acquisition.

In this study, expression of SSV2 Cas4 strongly reduce spacer acquisition, compared to the cells lacking SSV2 Cas4 (Fig. 7A and B). High-throughput sequencing data also supports that viral Cas4 hinders *de novo* spacer acquisition in *Sulfolobus* (Fig. 7B). Analysing the detected new spacers from viral Cas4 expression strain, we find SSV2 Cas4 has no effect on defining the 5’-CCN PAM or spacer length (Fig. 7B and C), but effect the 3’-A/G motif (Fig. 7B). Together, these results suggest that some invasive genetic elements may have accrued Cas4 proteins to corrupt CRISPR-Cas spacer acquisition by strongly reducing the spacer acquisition efficiency. In turn, the mobile genetic elements may have evolved to escape from CRISPR-Cas interference by the Cas4-disordered adaptation module, as also inferred from bioinformatic analyses (37).

Although expression of SSV2 Cas4 only reduced spacer acquisition efficiency, but does not affect integration orientation or spacer length (Fig. 7), it cannot exclude the effects of viral Cas4 on atypical spacer acquisition. For example, it is shown recently that a *Campylobacter* bacteriophage-encoded Cas4 homolog induces *de novo* spacer acquisition exclusively from host DNA (41). We extract all the new spacers with less than 3 mismatches to genomic DNA from the this study (41) and analyse the 5’- and 3’- motifs of the cognate protospacers. However, no conserved PAM or 3’ motifs are detected on both ends of the protospacers, indicating two possibilities: (1) the presence of DNA interference module counter-selects the functional genome-derived spacers with conserved PAM, and (2) the viral Cas4 produces defective spacer acquisition that would prevent CRISPR-Cas targeting of the genomic DNA (data not shown). Taking that only host DNA, but no viral DNA, is sampled into CRISPR array, both possibilities could be reasonable.

During viral infections, archaeal and bacterial cells have also evolved counteracting defence mechanisms including the production of antisense RNAs. Thus, a high level of antisense transcripts encompassing *cas2* and the upstream region of *cas4* is detected upon STSV2 viral infection in *S. islandicus*. The antisense RNA yields increases significantly at around 7 days post infection and then gradually decreases (44). This raises the possibility that host cells employ antisense RNAs to control the Cas4 expression levels in order to minimise its influence on acquisition efficiency and the consequent inactivation of CRISPR-Cas interference against invasive genetic elements.

## MATERIAL AND METHODS

### Strains, growth, and transformation of *Sulfolobus*

*S. islandicus* strains employed, including the genetic host E233S (Δ*pyrEF*Δ*lacS)* (45), the E233S derivative cas-deletion mutants, and the overexpression strains, were cultured in SCVy medium at 78°C. *S. islandicus* cells were transformed by electroporation and transformants were selected on two-layer phytal gel plates as described earlier (45). Interference and control plasmid transformation efficiencies were calculated as cfu/μg DNA of constructed plasmids. For experiments measuring new spacer uptake at the leader-repeat region of CRISPR loci, cells were grown in SCVy medium.

### Plasmid construction

The *csa1*, *cas4*, *cas4*-mutants and viral *cas4* overexpression plasmids were constructed by cloning the *cas4* and its mutant genes into the pCsa3a plasmid (9) to form an operon with the *csa3a* gene. The *csa1* (SiRe_0760), *cas4* gene (SiRe_0763) and viral *cas4* (p22) gene were amplified from *S. islandicus* REY15A or the fusellovirus SSV2 by PCR using FastPfu DNA polymerase (TransGene, Beijing China) with the primer sets listed in Table S1, and cloned into *csa3a* overexpression plasmid pCsa3a (9), yielding pCsa3a-Csa1, pCsa3a-Cas4 and pCsa3a-vCas4 plasmids, on which additional Shine-Dalgarno sequences were generated for translational initiation of Csa1 and host and viral Cas4 proteins. For mutations in the *cas4* gene, linear plasmids which were amplified from pCsa3a-Cas4 by PCR using FastPfu DNA polymerase contained the designed *cas4* gene mutations. After purification, these linear plasmids were cyclized by Gibson assembly before transforming into *E. coli* BL21 cells.

To construct interference plasmids for testing the flipped spacers, S10 spacer of CRISPR locus 2 with different 5’-end PAM and 3’-end sequences (Fig. 5A) were cloned into the pSeSD vector. For testing the slipped spacers, a protospacer sequence from the *cmr2a* gene template strand and its “slipped” derivatives were cloned into the site between two repeat sequences to form the mini-CRISPR under the control of the *araS* promoter in pSeSD. Interference plasmids were electroporated into *S. islandicus* strain E233S and transformation efficiencies were calculated. Primers used for constructing overexpression and interference plasmids are listed in Supplementary Table 2.

### PCR amplification of the integrated spacers

Transformants were cultured in 10 ml SCVy medium at 78°C until OD_600_ reached 0.3. Samples were taken from each culture (0.1 ml), and DNA was extracted from cells and employed as a PCR template. Leader-proximal regions of CRISPR locus 1 were amplified using Taq polymerase with forward and reverse primers CRISPR-F2 and CRISPR1S2-R (or CRISPR1S5-R). PCR products were separated by 1.5% agarose gel electrophoresis and visualised by ethidium bromide staining to identify the expanded bands. The PCR products were purified using PCR clean-up kit and sent for high-throughput sequencing. Primers used to amplify the leader proximal regions are listed in Supplementary Table 1.

### High-throughput sequencing and bioinformatics analysis

Six single colonies on each transformant plate were picked up and mixed in a single tube containing SCVy medium for growth. Genomic DNA was extracted from each transformant and used for amplification of the leader-proximal region. The leader-prxomal regions were also amplified from two samples of single colonies carrying *csa3a* and viral *cas4* overexpressing plasmid. Same amounts of the purified leader-proximal PCR products were subjected to HiSeq3000 sequencing (National Key Laboratory of Crop Genetic Improvement). After pair-end data assembly and low-quality data filtration, reads containing multiple (≥2) repeats were selected. For reads containing two repeats, the intervening sequence was considered as the original first spacer (S+1). For reads with three or four repeats, the leader-proximal spacers (S-2 and S-1) were considered as the new spacers. Using the BLASTN program against pSeSD plasmid (35) or the *S. islandicus* REY15A genome sequence (31), the protospacer sequence was identified for each spacer. The 3-bp sequence at 5’-end of each protospacer was considered as the PAM region. Perl scripts were run to analyse the protospacers. All high-throughput sequencing reads are deposited at the SRA database with the accession number PRJNA498890. The conserved motifs of the 5’-end and 3’-end adjacent sequences of the protospacers were analysed using the WebLogo (46). The neighbour-joining tree was generated from a T-coffee alignment of the proteins using Mega7 (47, 48), with pairwise distances between sequences uncorrected.

## ACKNOWLEDGEMENTS

We thank Roger A Garrett and Qunxin She of the Danish Archaea Centre in the University of Copenhagen, Denmark, for critical reading of the manuscript. This work was supported by the National Natural Science Foundation of China (No. 31671291 to N.P.), the National Postdoctoral Program for Innovative Talents (No. BX20180112 to T.L.), and the Fundamental Research Funds for the Central Universities (No. 2662015PY038 and 2662015PX199 to N.P.). Funding for open access charge: the National Natural Science Foundation of China (No. 31671291).

